# Combined effects of P25 TiO_2_ nanoparticles and disposable face mask leachate on microalgae *Scenedesmus obliquus*: Analysing the effects of heavy metals

**DOI:** 10.1101/2023.06.08.544143

**Authors:** Soupam Das, Amitava Mukherjee

**Author notes:** Corresponding Author Amitava Mukherjee Senior Professor and Director Centre for Nanobiotechnology Vellore Institute of technology, Vellore 632014, India.

## Abstract

Disposable surgical masks have been extensively employed as protective medical equipment due to the widespread breakout and transmission of the COVID-19 virus across the globe. These masks were made up of plastic polymer materials that would emit microplastics after entering the environment. Therefore, their careless disposal might lead to new and bigger microplastic contamination. The impacts of plastics that seep into waterways and their subsequent interactions with aquatic life are yet largely unexplored. In this study, we determined the quantity and kind of microplastics that were discharged from disposable surgical face masks. Furthermore, we also quantified heavy metals leached from the face masks (HML). In contrast, the increasing usage of nTiO_2_ in consumer items has led to its ubiquitous presence in freshwater systems. Four different concentrations of nTiO_2_, 0.5, 1, 2, and 4 mg L^-1^ were mixed with face mask leachates (FML) to perform the mixture toxicity test on freshwater algae, *Scenedesmus obliquus*. Reduced cell viability and photosynthetic activity were noticed in the treatment groups containing nTiO_2_ and FML. This was accompanied by increased oxidative stress and antioxidant activities. Furthermore, the heavy metals leached from the face masks were also tested for toxicity. In addition to that, changes in the cellular morphology were also studied with the help of FE-SEM and FTIR analysis. Our study reveals that leachates from disposable surgical face masks along with nTiO_2_ possess a serious threat to the environment.

**Environmental significance:** During COVID-19, surgical face masks were widely used and discarded. These discarded face masks end up in lakes, rivers, and ponds. The facemasks were composed of polypropylene and other polymers. These masks release microplastics and heavy metals when discarded into water bodies. The current research focuses on assessing the environmental toxicity of the microplastics and heavy metals leached from the masks using algae as a model system. Our work further demonstrates the combined toxic effects of nTiO_2_ in the presence of the face mask leachate. Algae plays a crucial role as the primary producer in the freshwater ecosystem. These emerging contaminants may act as environmental stressors to the microalgae, and this may impair the ecosystem’s structure and function.

## 1. Introduction

The COVID-19 pandemic has expanded the global usage of disposable face masks (DFMs) to protect the health of human beings ^1^. A face mask prevents the spread of germs via contact with body fluids and vice versa ^2^. Billions of face masks are worn daily across the globe, and many of these masks eventually make their way into freshwater systems via various water bodies ^3^. These disposable surgical face masks are made from plastic polymers such as polyethylene and polypropylene (PP) ^4^. The ubiquitous and inevitable usage of face masks has caused serious worries over the development of enormous quantities of plastic trash. This enormous amount of trash has the potential to become a significant contributor to environmental microplastics ^5, 6^. Microplastics may be released from disposable face masks by regular wear and tear or by natural weathering processes like chemical oxidation, UV degradation or changes in temperature and humidity ^7, 8^.

Primary producers like microalgae support the freshwater habitats ^9^. A significant number of research has evaluated the toxic effects of microplastics on various freshwater microalgae. Xiao and his colleagues observed dose-dependent decrease in growth rate of freshwater microalgae *Euglena gracilis* upon exposure to increasing concentrations of microplastics ^10^. Similarly, Zheng et.al.^11^ observed reduced growth and development of *Microcystis aeruginosa* when interacted with 60 nm microplastics in a dose-dependent manner.

Engineered nanoparticles have also been recognized as potential new pollutants in a freshwater ecosystem ^12^. Titanium dioxide nanoparticles (nTiO2) were found to be highly toxic to algae, plants and fishes ^13^. TiO2, being a semiconducting material, can absorb UV light at wavelengths that are shorter than its band gap, which results in the production of electron–hole pairs and the activation of photocatalytic processes. nTiO2 have a greater number of atoms on their surfaces than their bigger (micron-sized or bulk) counterparts. For these reasons, nTiO2 is utilized in a wide variety of consumer goods, including solar panels, sunscreen, self-cleaning coatings, germicidal sprays, and water purifiers ^14^. Therefore, an uncontrolled release of nTiO_2_ into the aquatic environment may take place, which may result in detrimental impacts on aquatic life, particularly algae, which are the main producers in aquatic trophic chains. One of our earlier lab research found that UV-B irradiated nTiO_2_ was more toxic than UV-A towards freshwater *Chlorella sp* ^15^. The cytotoxicity of the freshwater algae *Chlorella pyrenoidosa* was investigated by researchers, who observed that the particle size and dispersibility of nTiO2 have an important influence on the cytotoxicity of the organism ^16^. According to the findings of certain research, nTiO_2_ inhibited the development of algae and reduced the amount of total chlorophyll content ^17^.

At this point, the majority of the published data are related to the risk evaluation of individual microplastics as well as pristine nTiO_2_ on microalgae. There have only been a handful of studies that examined the combined toxicity of leachates from the face mask and nTiO_2_ in algae. Huang and his research group investigated the combined toxic effects of silver nanoparticles and nanoplastics on *Chlamydomonas reinhardtii* and *Ochromonas Danica*. They observed that nanoplastics enhanced the toxic effects of silver nanoparticles towards both the freshwater algae ^18^. Similarly, Bellingeri et. al.^19^ conducted research on the toxic effects of copper and Carboxylated nanoplastics towards *Raphidocelis subcapitata*. There was no significant difference in the rate of growth inhibition that was detected between exposure to copper alone and exposure to copper in combination with nanoplastics.

There are only a few research papers dealing with the potential ecotoxicological effect of plastics and other heavy metals leached from face masks ^20, 21^. This is the first study to investigate the potential role of leachates from single-use face masks in amplifying the toxicity of nTiO_2_ in freshwater algae *Scenedesmus obliquus*. The objectives of this study are (i) isolation and quantification of microplastics and heavy metals from disposable surgical face masks (ii) toxicity study of mask leachate on *Scenedesmus obliquus* (iii) toxicity study of mask leachate in combination with nTiO_2_ on algae. Additionally, changes in the cell morphology after exposure to toxicants were also studied.

## 2. Materials and methods

### 2.1. Materials

We bought chemicals like hydroxylamine hydrochloride, dimethyl sulfoxide (DMSO), thiobarbituric acid (TBA) and trichloroacetic acid (TCA) from Hi-Media Pvt. Ltd (Mumbai, India). Aeroxide P25 nTiO_2_ (Titanium (IV) dioxide nanoparticles), 2’,7’ dichlorofluorescein diacetate (DCFH-DA) were procured from Sigma-Aldrich (USA). We purchased the nitro blue tetrazolium (NBT) dye and the hydrogen peroxide solution (H_2_O_2_-30% w/v) from SDFCL in Mumbai, India.

The surgical face masks were acquired from local medicine shops and pharmacies. All masks were within their expiry date and there are no signs of damage. The masks have been measured to be 17 x 9.5 cm. The packaging indicates that there are three distinct layers to the mask. The outside layer is produced from a non-woven fabric that is leak-proof, the layer in the centre is made from high-density filter ply, and the lower layer is created from soft fabric. The colour of the outermost layer is a pale blue, but the colour of the middle and the inner is white, as shown in Fig. S1.

The interaction medium used for this research was freshwater to imitate natural environmental conditions. The source of freshwater was the lake located inside the campus of Vellore Institute of Technology, Vellore, India (12.9692° N, 79.1559° E). To eliminate bigger particles from the collected lake water, it was filtered many times using blotting paper and with filter paper of pore size 11 µm (Whatman No.1). Following filtration, it was then autoclaved at 121 °C for 15 mins at 15 lbs pressure. It then underwent a second round of aseptic filtering using a 0.1-micron filter. Any further filtering techniques had to be avoided so that its physiochemical qualities could be preserved. In prior investigations, we provided the complete characterization of lake water ^22^. All of the experiments conducted here utilized this sterilized lake water as their matrix ^15^.

### 2.2. Preparation and characterization of nTiO2 stock suspension

To determine the surface characteristics of the nanoparticle, nTiO2 powder was subjected to a field emission scanning electron microscope (FE-SEM) after sputter coated with gold.

The stock solution (100 mg L^-1^) of nTiO_2_ was prepared in Milli-Q water. For proper dispersion of the nanoparticle, an ultrasonicator (Sonics, USA; 130 W, 20 kHz) was employed to sonicate the solution for 40 mins under dark conditions.

### 2.3. Microplastic analysis

The mask borders, metal nose piece and elastic ear loops were snipped off using a pair of scissors. The full face masks were then placed in a 500 ml screw-top glass container containing 200 ml of sterile lake water. The containers were shaken at 120 rpm for 24 h on a rotary shaker. A bottle containing only water served as control. After 24 h, the leachate solutions were transferred into a clean glass bottle. Three replicates were used for each treatment ^4, 23^.

### 2.4. Quantification, identification and characterization of the leached microplastics

Using a field emission-scanning electron microscope (FE-SEM) (Thermo Fisher FEI Quanta 250 FEG), the number of microplastics discharged from the leachate were measured and counted. 100 µl of the leachate solution was dropped on a glass slide and allowed to dry at room temperature ^24^. The glass slide was then sputtered with gold and analysed under FE-SEM. Image J plus software was used to count the number of particles from the FE-SEM images. For each face mask and its replicates, twenty photos were picked at random for analysis ^4^.

Additional characterization of the mask leachates was achieved via FTIR analysis, following the previously published procedure ^4^. A cellulose membrane filter (0.8 µm pore size) was used to filter the mask’s leachates. To ensure that the samples were thoroughly dry, they were freeze-dried and then lyophilized while still in the membrane filter. FTIR (IR Spirit; Shimadzu, Japan) was used to characterize the dried samples and compare them to reference plastic standards.

Raman spectroscopy was performed to verify the released particles as microplastics. The leachates were filtered through a cellulose membrane filter and kept for drying on a glass Petri plate at room temperature. The membrane filter containing the dried microplastic fibres was analyzed by Raman spectroscopy (Anton Paar, Cora 5001, Austria). Raman spectra were acquired using a laser excitation wavelength of 785 nm. The range of the spectra was from 300 to 3200 cm^-1^. The spectra of the samples were entered into the Polymeric Materials Database, and the kinds of polymers were identified based on the degree to which their spectra were similar to those of the standards ^7^.

Inductively coupled plasma-optical emission spectrometry (ICP-OES) was employed to analyse and quantify any heavy metals present in face mask leachate. The leachate solution was filtered using 0.45-micron filter paper to separate the microplastic from the leachate solution. The filtered leachate solution was then analysed by ICP-OES.

### 2.5. Test organisms

In this study, a freshwater microalga *Scenedesmus obliquus* was employed. These algae were obtained from the lake at Vellore Institute of Technology (VIT) by using a previously described process ^17^. In a sterile BG11 medium, the isolated algal cultures were maintained and occasionally subcultured. For the optimal development of the algae, the cultures were maintained in cycles of light and dark. For illumination, a 3000 lux fluorescent bulb was employed (Philips TL-D Super 80 Linear fluorescent lamp, India). During this whole study, the OECD guidelines were followed strictly ^25^.

### 2.6. Toxicity studies

#### 2.6.1 Experimental setup

For this experiment, 15-day-old (exponential phase) algal cells were taken and centrifuged at 7000 rpm for 10 mins at 4 °C. The pellet was rinsed twice with sterile lake water to remove any residual growth media. We obtained an algal cell density of 0.5 OD (4 x 10^3^ cells per mL) by diluting the cell pellets with sterile lake water medium, which was then treated with different concentrations of nTiO_2_ (0.5, 1, 2, and 4 mg L^-1^) and 5.4 x 10^3^ particles/ml of FML in a 50 ml glass beaker. The EC_50_ (1.3 mg L^-1^) of nTiO_2_ from our prior investigation was used to determine the working concentrations of nTiO_2_ ^15^. We picked two concentrations that were lower than the EC_50_ value and two values that were higher than the EC_50_ value. The concentrations of the leachate from the mask were determined to be 5.4 x 10^3^ particles/ml, as described in the results section. Each concentration of nTiO_2_ was mixed with 5.4 x 10^3^ particles/ml of FML and incubated for 72 h under visible light conditions. The algal suspensions that were not treated with nTiO_2_ or FML served as a control for the experiment. In addition, microplastics were separated from the leachate solution using 3KDa filter paper. Algal cells were then treated with filtered leachate solution containing heavy metals. Natural organics in the lake water cannot affect either the treatment group or the control group since both are incubated in the same lake water matrix. To avoid the cells from adhering to the bottom or the sides of the beaker, the samples were shaken once a day. To determine statistical significance, each study was conducted in triplicates (n = 3).

#### 2.6.2. Determination of algal cell viability

After 72 h incubation, 10 µl of the samples were placed on a Neubauer chamber and observed under an optical microscope to count the number of viable cells (Leica DM2500). The percentage of algal cell viability was calculated relative to the control. To compute the percentage of viable cells, the following formula is used:

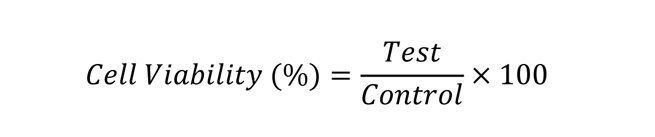

### 2.7. Morphological changes in Scenedesmus obliquus

FE-SEM analysis was performed to observe the changes in size and shape of *Scenedesmus obliquus* after treatment with nTiO_2_, FML and their binary mixture. After 72 h, a drop of the incubated cell solution was placed on a glass slide and air-dried. The slide was then sputter coated with gold and examined under an electron microscope to observe the changes in algal cells.

### 2.8. Analysis of oxidative stress

The fluorescent dye DCFH-DA was used to calculate the total reactive oxygen species (ROS) generated by the treated algal cells. After the algae had been exposed to the toxicants for 72 h, samples were incubated with DCFH-DA dye (50 µL of 100 µM) for 30 mins in the dark. After incubation, the fluorescence intensity of the samples was evaluated using a fluorescence spectrophotometer (Cary Eclipse, G9800A; Agilent Technologies, USA). The excitation and emission wavelength were at 485 nm and 530 nm respectively ^26^. The obtained intensities were compared to the intensities of the control algal cells. Under abiotic conditions, there was no apparent increase in the fluorescence intensity.

The MDA concentration was determined by subjecting the interacted algal samples to 30 mins of heating in a water bath at a temperature of 95 °C along with the mixture of TBA and TCA (0.25% (w/v) TCA in 10% (w/v) TBA). Following centrifugation, a UV-Vis spectrophotometer (xMARK microplate absorbance spectrophotometer, BIO-RAD) was used to evaluate the absorbance of the supernatants at 532 and 600 nm ^27^.

### 2.9. Analysis of antioxidant activity

The comprehensive method for analyzing superoxide dismutase (SOD) activity can be followed from the study published by Kono in 1978 ^28^. After incubation, the cell pellet was recovered by centrifuging the sample at 7000 rpm for 10 mins at 4 °C, washed, and then homogenising it in 0.5 M phosphate buffer before centrifuging it again for 20 mins at 4 °C. Following centrifugation, the following components were added to 100 µl of the supernatant: 50 mM Na_2_CO_3_, 96 mM nitro tetrazolium blue chloride, 0.6% triton x-100, and 20 mM hydroxylamine hydrochloride. This mixture was then incubated at 37 °C for 20 mins under visible light conditions. Following incubation, the absorbance of these samples was determined using a UV-spectrophotometer (xMARK microplate absorbance spectrophotometer, BIO-RAD) at 560 nm.

To perform the catalase assay, the interacted algal samples were centrifuged at 7000 rpm for 10 mins at 4 °C after being incubated for 72 h. One of the past study on catalase assay showed the detailed procedure to perform this experiment ^29^. After centrifugation, they were rinsed with sterile 0.5 M PBS and homogenized. After adding 1 ml of a newly prepared H_2_O_2_ solution (10 mM) to 2 ml of supernatant, the absorbance at 240 nm was measured for 3 minutes using a spectrophotometer (Hitachi U-2910 Japan). The reaction mixture without H_2_O_2_ was used as a control.

### 2.10. Analysis of photosynthetic yield

A photosynthesis yield analyzer (Mini PAM, made by Heinz Walz in Germany) was used to detect chlorophyll fluorescence as well as the electron transport rate in the samples. This was performed to quantify the level of stress that the toxicant was putting on algal photosystem II.

After 72 h of exposure, the samples were put in a dark condition for 30 mins. Following that, 200 µl of the interacted algal samples were placed into the chamber of the Photosynthesis Yield Analyzer (Mini PAM), and a high-intensity actinic light was used to evaluate the Fv/Fm ratio and the ETR max value. The ratio of the maximum photochemical quantum yield of the PS II system in treated and control algal cells is denoted by the symbol Fv/Fm, where Fv stands for variable fluorescence and Fm stands for the maximum fluorescence released by the samples when exposed to high intensities of actinic light. Graphs were used to illustrate these findings in comparison to the samples used as controls.

### 2.11. Surface functional group analysis using FTIR

FTIR was utilized to understand more about the role of surface functional groups in the reaction between algae, nTiO_2_, FML and their binary mixture. After the interaction, the algal cells were centrifuged at 10,000 rpm for 15 mins. The pellet was collected and freeze-dried for 24 h in a (-80) refrigerator. The frozen samples were kept for lyophilisation. The FTIR spectrum was analyzed from lyophilized algal pellets (FTIR make: IR Spirit, Shimadzu, Japan).

### 2.12. Quality assurance and control

Every piece of equipment used for sampling or analysis in the lab was cleaned with sterile DI water and then wrapped in aluminium foil to avoid cross-contamination. During the sampling and examination procedure, clean gloves, cotton masks, and cotton lab coats were used. Glassware was pre-cleaned with an ethanol solution, washed extensively with Milli-Q water (pH 7.0), and then heat-treated at 400 °C to eliminate any organic contaminants. All solvents used were of analytical quality (>95%). To minimize microplastic contamination, metal or glass items were used. Additionally, the researcher avoided interaction with products that were comparable to the mask material.

### 2.13. Statistical analysis

For this study, each experiment was performed three times (n=3). Each data set is represented by the mean standard deviation. Using Graph-pad Prism V8.0 software, a two-way ANOVA (Bonferroni) was performed to determine the statistical significance of the data.

## 3. Results and discussion

### 3.1. Characterization of nTiO_2_ and FML

Fig. 1A represents FE-SEM analysis of nTiO_2_ which shows spherical or cuboidal-shaped particles with a size of 20 nm. Microplastics released from the face mask were irregular in form, with sizes ranging from 1µm to 15 µm, as shown in FE-SEM images (Fig. 1B). In addition, the FE-SEM data revealed that the mask released 2.88 x 10^6^ particles/L. Disposable surgical face masks of all brands shed large amounts of irregularly shaped microplastics. During the SEM study, only a small amount (100 µL) of leachate was analysed, thus it is hard to say for sure whether or not the mask includes bigger bits of plastic debris. For instance, researchers have observed that standard masks may include large plastic fibres of 600 to 1800 µm in diameter ^30^.

**Fig. 1:**
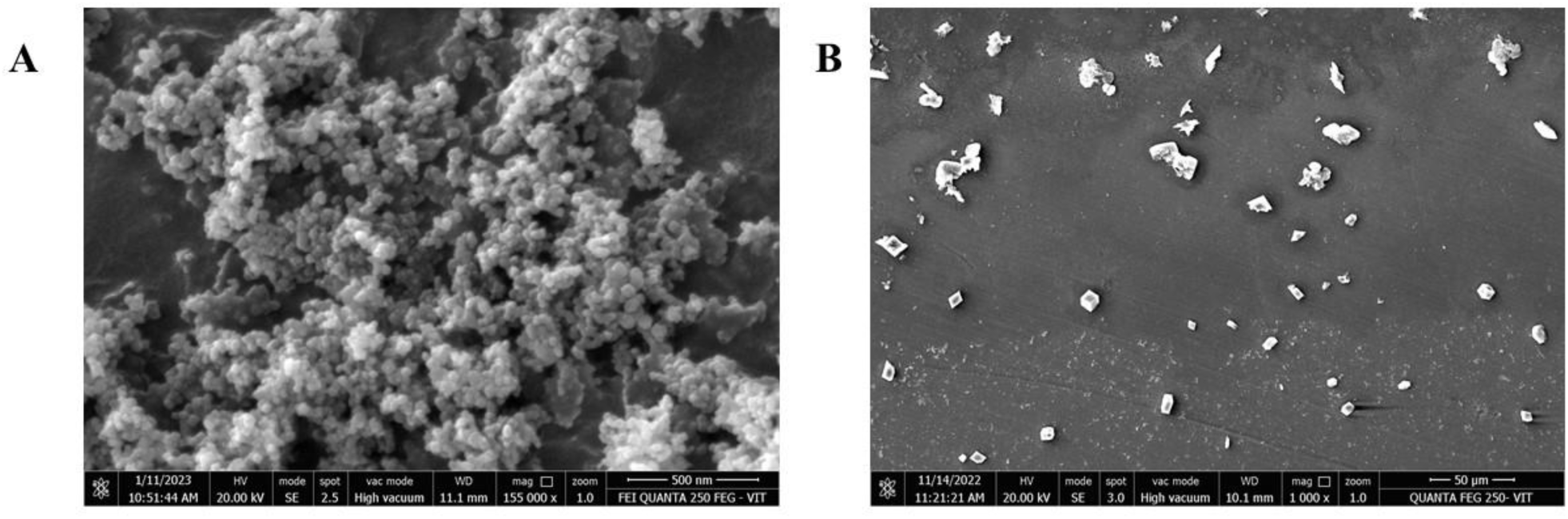
Field emission scanning electron microscopic (FE-SEM) images of (**A**) nTiO_2_ (**B**) FML

The FTIR spectra of the microplastics in FML are shown in Fig. 2. The spectral bands of particles leached by the mask were studied, and the results exhibited specific bands indicative of Polypropylene. The Raman spectra indicate the presence of microplastics in the mask leachates (Fig. S2). It consisted of 85% polypropylene and other fibres. Similar FTIR and Raman characteristics were observed in our previous investigations on FML ^31^. Disposable facial masks were found to have leached heavy metals when the FML was analysed by ICP-OES. The heavy metals and their concentrations in FML are shown in Table S1.

**Fig. 2:**
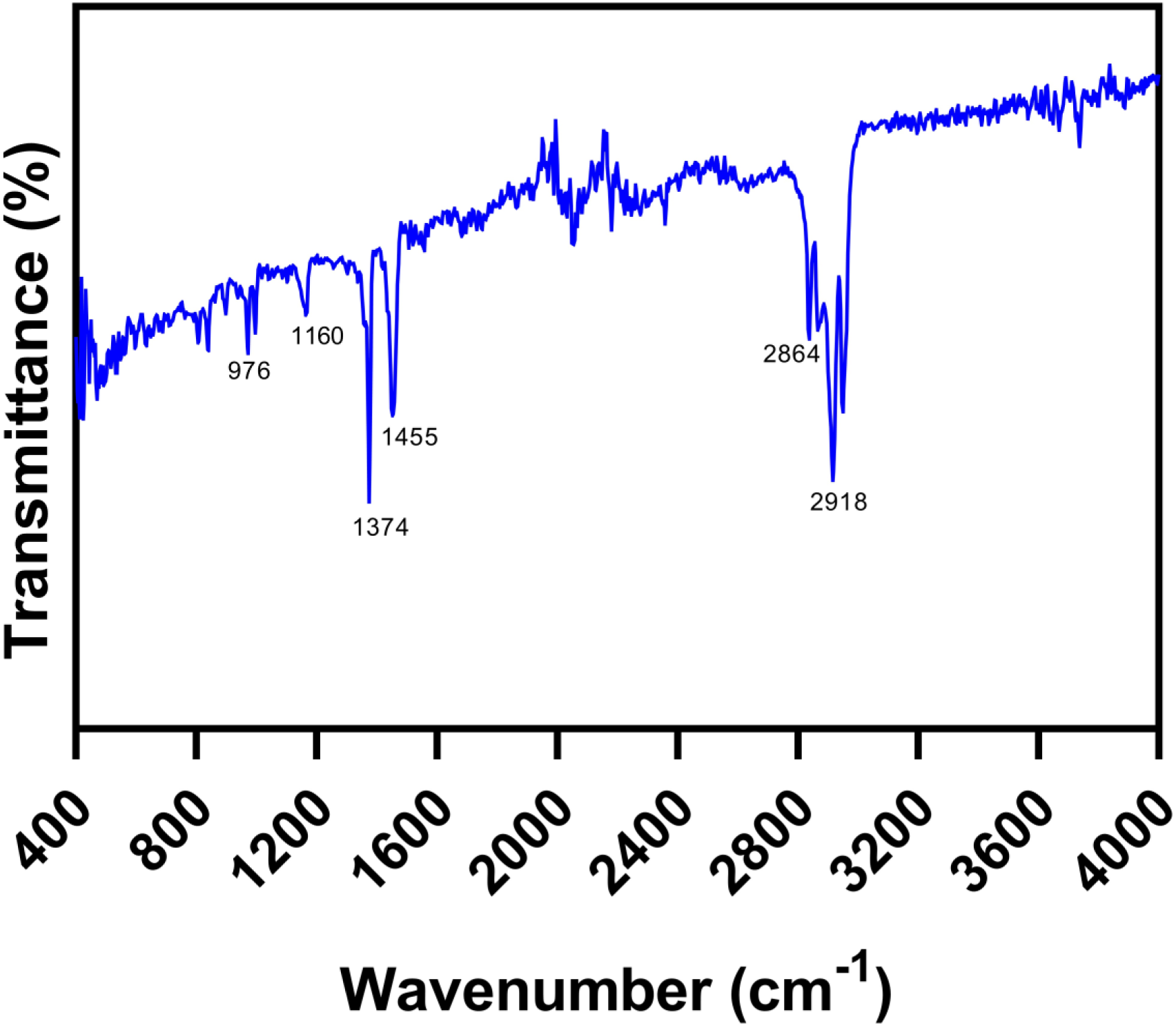
FTIR spectra of disposable surgical face mask leachate (FML)

### 3.2. Influence of FML on the toxic effects of nTiO_2_

*Scenedesmus obliquus* treated with nTiO_2_, FML, and their binary combination exhibited a dose-dependent significant reduction in cell viability, as shown in Fig. 3A (p <0.001). The reduction in cell viability was found to be highly significant compared to the control for all the treatment groups (p<0.001). Furthermore, cells treated with a binary mixture also showed reduced viability when compared with pristine nTiO2. In addition, algal cells when treated with HML showed a toxicity of 5.9% (Fig. S3A).

**Fig. 3:**
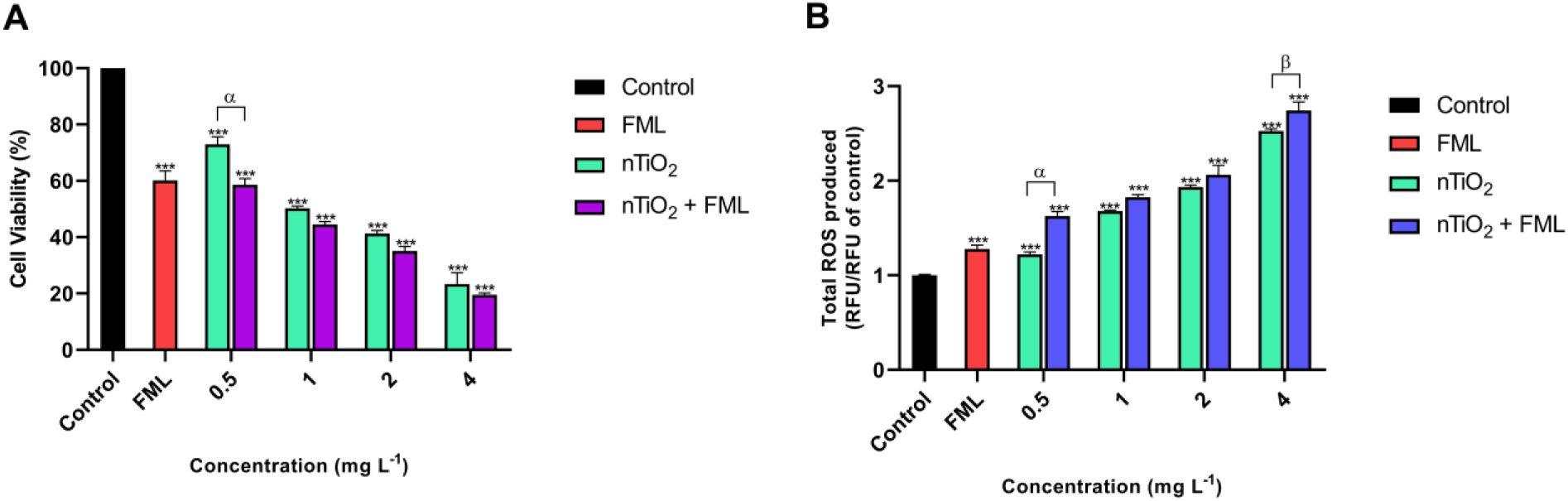
**(A)**Decrease in cell viability in percentage when interacted with nTiO_2_, FML and their binary mixture. **(B)** Total ROS produced when interacted with nTiO_2_, FML and their binary mixture. ‘*’ symbolizes a significant difference in percentage observed with respect to control (*** = p < 0.001). ‘α, β’ symbolizes a significant difference observed between the treatment groups (α = p < 0.001, β = p < 0.01)

The adsorption of microplastics onto the surface of algal cells may hinder energy transmission and cause stress, especially if the microplastics are of a larger size ^32^. The previous findings corroborated the toxicity results presented here, showing that microplastics reduce growth rates regardless of their size, concentration, or type ^33, 34^. Based on the findings of this study, it is possible to hypothesize the mechanism by which nTiO_2_ is toxic to *Scenedesmus obliquus*. Previous studies on nTiO_2_ toxicity towards *Scenedesmus obliquus* also showed similar results ^35^. After interaction with HML, much decrease in cell viability was not observed. The major reason for this is that masks leached very low levels of heavy metals. It could enhance oxidative stress but does not cause any increased cell death. Previous research showed that with rising concentrations of heavy metals, toxicity on microalgae also increased ^36^.

### 3.3. Morphological changes in Scenedesmus obliquus

After 72 h of interaction, there was no indication of algal cell death or deformation in the control group (Fig. 4A). Whereas, when algal cells were treated with pristine nTiO_2_ (4 mg L^-1^), FML and their binary mixture, cell deformity and cell death were observed. In addition, FE-SEM images also revealed that both nTiO_2_ and microplastics were adsorbed onto the surfaces of the algal cells as evident from Fig. 4B, C, & D.

**Fig. 4:**
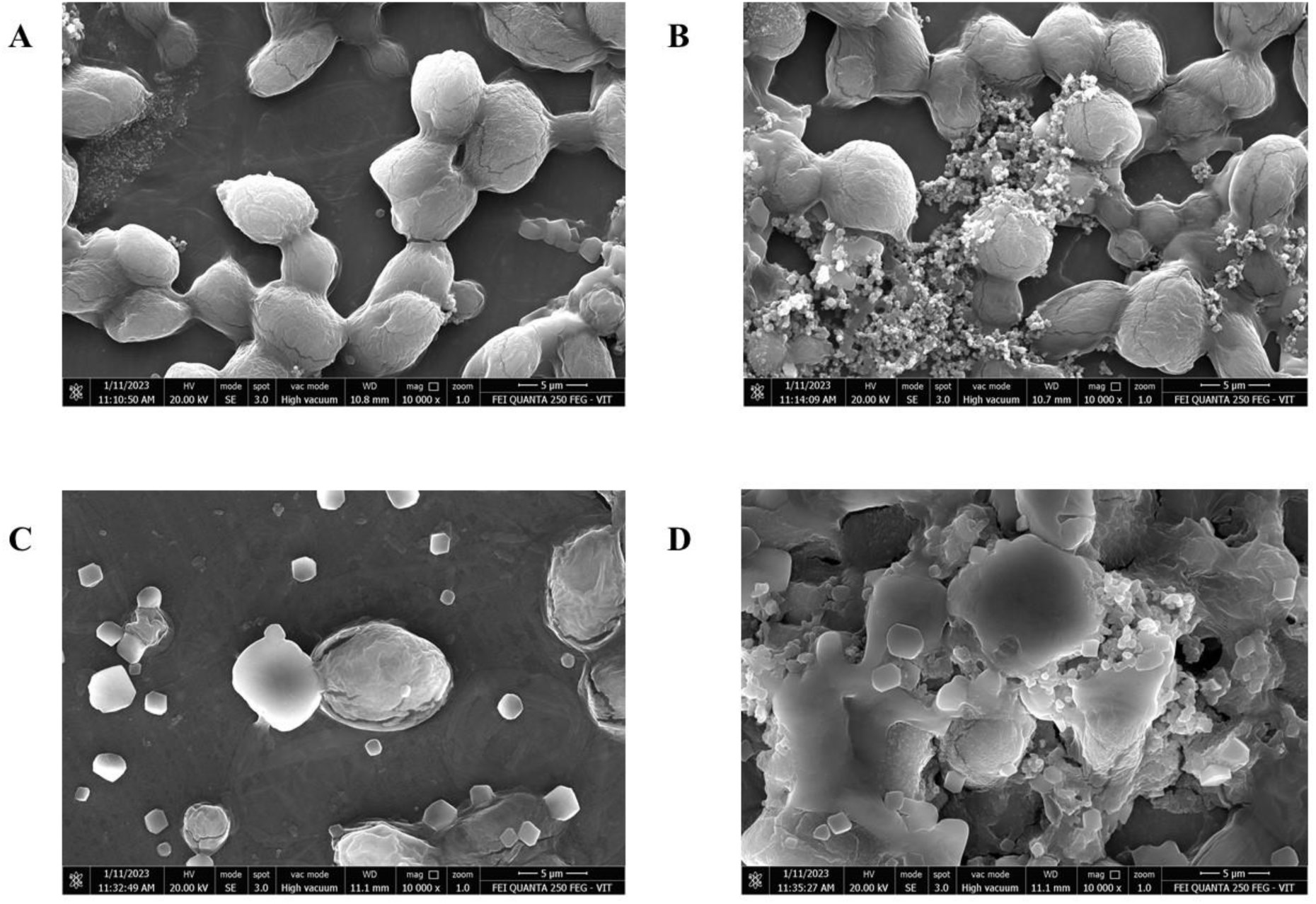
Morphological changes of *Scenedesmus obliquus* observed under FE-SEM after 72 h of incubation **(A)** Control **(B)** algae exposed to nTiO_2_ (4 mgL^-1^) **(C)** algae exposed to FML **(D)** algae exposed to nTiO_2_ (3.2 mgL^-1^) + FML

In Fig. 4B it is observed that nTiO_2_ was adsorbed onto the algal cell surface. nTiO_2_ adsorption on the cell surface may cause physical and chemical disruption of cellular activity^37^. Physically when the particles were adsorbed onto the algal cells, they impede the movement of electrolytes and metabolites across the cell membrane ^38^. Irradiating nTiO_2_ with light at a wavelength less than that of the band gap energy will excite the excitons, resulting in holes being created in the valence band. This occurs on a chemical level since nTiO_2_ is a photocatalyst. Both the electrons and the holes are powerful oxidizing and reducing agents, respectively. Free reactive oxygen species radicals are produced when electrons and holes react with water molecules ^37^.

### 3.4. Oxidative stress generation

Fig. 3B represents the total ROS produced by the algal cells after interacting with nTiO_2_, FML and their binary mixture. A dose-dependent increase in ROS generation was observed when the algal cells were treated with nTiO_2_ alone, FML alone and their binary mixture. The increase in ROS generation for all the treatment groups was statistically significant as compared to the control (p<0.001) The amount of ROS generated by the binary mixture treated algal cells was statistically significant compared to 0.5 and 4 mg L^-1^ of nTiO_2_ alone respectively. A minor increase in ROS was observed when the algae were treated with HML (Fig. S3B). Statistically significant (p<0.001) increases in LPO generation were detected across all concentrations of nTiO_2_, FML, and their binary mixture as compared to the control (Fig. 5A). The difference observed in LPO generation between the binary mixture and their corresponding nTiO_2_ concentrations was also found to be statistically significant (p<0.01) except for 2 mg L^-1^ of nTiO_2_. Similar to ROS results, a minor increase in LPO was observed when the algal cells were treated with HML (Fig. S4A).

**Fig. 5:**
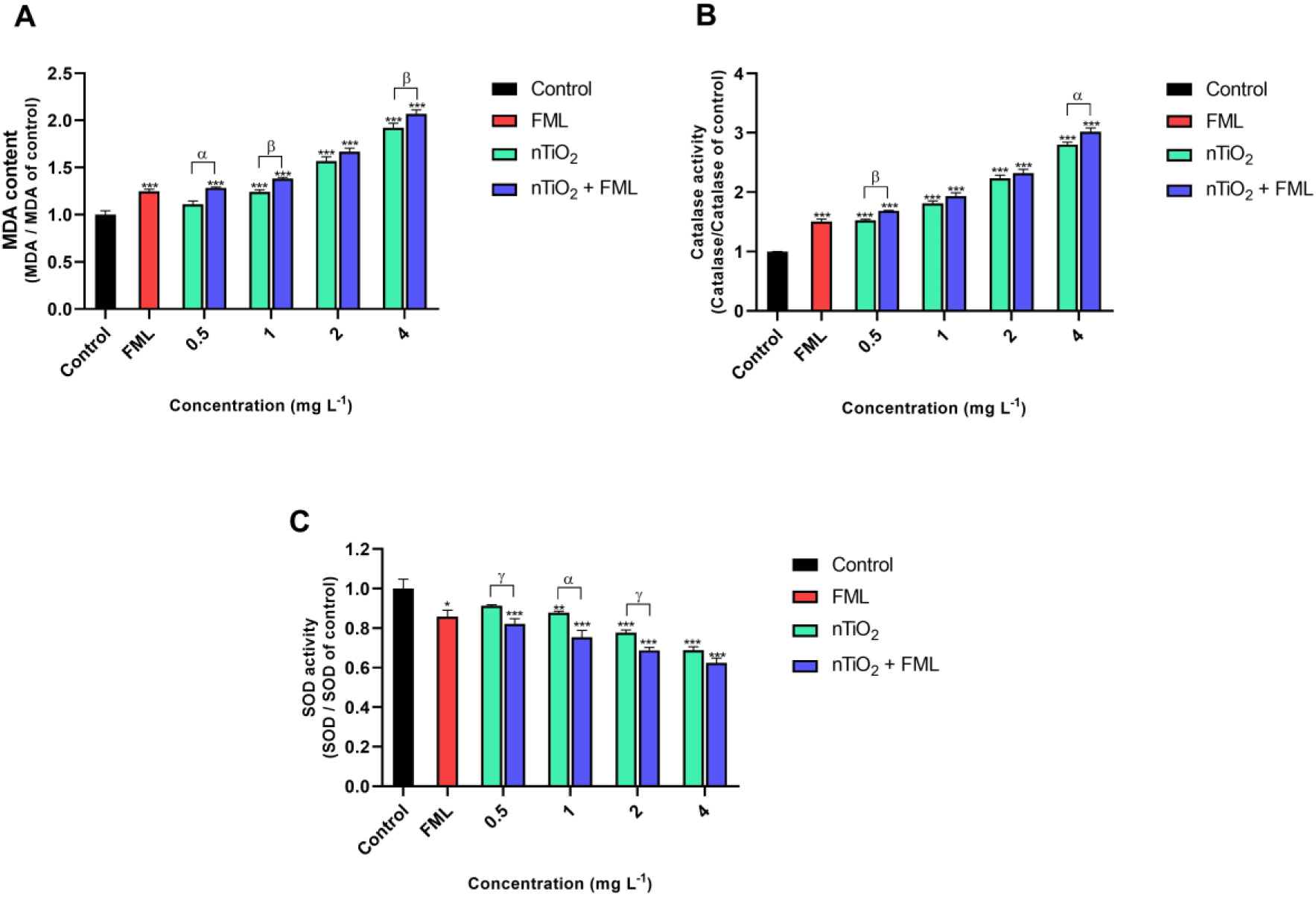
**(A)** LPO produced when interacted with nTiO_2_, pristine FML and their binary mixture. (**B**) CAT activity when interacted with nTiO_2_, FML and their binary mixture (**C**) SOD activity when interacted with nTiO_2_, FML and their binary mixture. ‘*’ symbolizes a significant difference in percentage observed with respect to control (*** = p < 0.001). ‘α, β, γ’ symbolizes a significant difference observed between the treatment groups (α = p < 0.001, β = p < 0.01, γ = p < 0.05)

From the results of ROS and LPO production, a direct correlation can be established between the production of oxidative stress and the cell viability of *Scenedesmus obliquus*. This could lead to membrane damage since the reactive oxidant species might burn the fatty acids found in the membrane phospholipids. This explains why ROS generation and lipid peroxidation are positively correlated ^39^. A recent research showed enhanced ROS and LPO generation in *Chlorella sp*. and *Raphidocelis subcapitata* when exposed to copper and nanoplastics mixture ^40^. As previously mentioned, *Scenedesmus obliquus* generated a higher level of ROS, which increased the amount of damage to the cell membrane ^41^. The results of our study are in agreement with our previous studies, which found that exposure to nano plastics and nTiO2 decreased the metabolic activity of microalgae *Scenedesmus obliquus* ^35^. HML caused a moderate increase in oxidative stress as discussed in the previous section. But with increased concentration of these heavy metals, it might affect the fatty acids on the cell membrane and can reduce cell viability ^42^. Moreover, these metals may indirectly promote oxidative stress by interfering with metabolic activities such as respiration and photosynthesis, depleting GSH and inhibiting antioxidant enzymes ^43^.

### 3.5. Antioxidant assays

A concentration-dependent increase in CAT activity was observed for nTiO2, mask leachates, and their binary mixture which is statistically significant to control (p<0.001). CAT activity increased significantly for the binary mixtures when compared to their pristine counterparts (except for 1 and 2 mg L^-1^ of nTiO_2_) (Fig. 5B). Furthermore, treatment of the algal cells with HML resulted in a small increase in CAT activity (Fig. S4B). Similarly, a concentration-dependent decrease in SOD activity was observed for nTiO_2_, mask leachates, and their binary mixture which is statistically significant to control (p<0.001). SOD activity decreased significantly for all the binary mixtures when compared to their respective single exposure of nTiO2, except for 4 mg L^-1^ of nTiO_2_ (Fig. 5C). When the algal cells were exposed to HML, a modest reduction in SOD activity was also detected (Fig. S4C).

Antioxidant enzymes are the countermeasures used to reduce the amount of oxidative stress. The reduced SOD activity in the algal cells may be an indication that the ROS generated is inhibiting the SOD enzyme ^44^. CAT converts hydrogen peroxide to water and oxygen, whereas SOD is efficient at neutralizing superoxide radicals. Increasing SOD and CAT levels may help to manage high levels of ROS ^45^. In contrast to the prior findings, increasing the concentration of nTiO2 alone or the binary combination led to a reduction in SOD while simultaneously leading to an increase in CAT. As a consequence of excessive ROS production, the SOD in algae may become active and convert the produced superoxide radicals into hydrogen peroxide. The disproportionation process generates H_2_O_2_, which, when combined with the H_2_O_2_ already present in the system, might potentially damage the SOD enzyme and bring the enzyme’s activity level down. Disproportionation generates H_2_O_2_, which combines with any pre-existing H2O2 to potentially damage the SOD enzyme, reducing its function ^35, 46^. *Scenedesmus obliquus* treated with nTiO_2_ and mask leachates showed elevated CAT activity. This suggests that intracellular H_2_O_2_ was removed by scavenging. Previous studies have demonstrated that microplastics may stimulate the production of anti-oxidant enzymes in *Microcystis aeruginosa* ^47^ and *Euglena gracilis* ^10^. HML increased CAT activity and lowered SOD activity significantly. This was a result of heightened oxidative stress. Cheng et. al. (2016) observed increased antioxidant activities of microalgae when treated with heavy metals like cadmium ^48^.

### 3.6. Maximum quantum yield of PSII and ETR

A significant decrease in photosynthetic activity compared to control was observed in *Scenedesmus obliquus* when interacted with nTiO_2_, FML and their binary mixtures (p<0.001) (Fig. 6A). Algal cells treated with binary mixer exhibited a moderate reduction in photosynthetic activity compared to pristine nTiO_2_ counterparts. Moreover, treatment of algal cells with HML resulted in a minor reduction in photosynthetic activity with respect to control (Fig. S5A). A significant decrease in Φm was observed in *Scenedesmus obliquus* when interacted with nTiO_2_, FML and their binary mixtures (p<0.001) compared to control (Fig. 6B). Compared to pristine nTiO_2_ counterparts, algal cells treated with binary mixer also exhibited a significant reduction in Φm. Furthermore, a negligible decrease in Φm was observed with respect to control after treatment of the algal cells with HML (Fig. S5B).

**Fig. 6:**
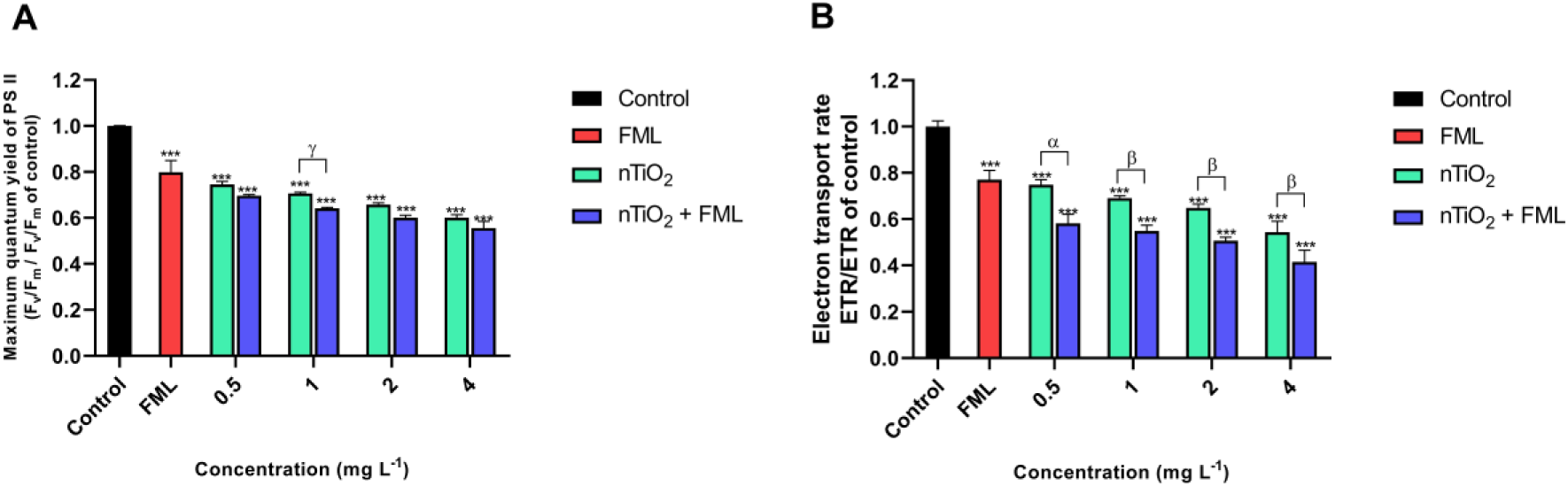
**(A)** Maximum quantum yield of PSII after exposure to nTiO_2_, FML and their binary mixture **(B)** Electron transport rate after exposure to nTiO_2_, FML and their binary mixture. ‘*’ symbolizes a significant difference in percentage observed with respect to control (*** = p < 0.001). ‘α, β’ symbolizes a significant difference observed between the treatment groups (α = p < 0.001, β = p < 0.01)

The mask-leached particles that were adsorbed on the surface of the algal cells also had a significant influence on the photosynthetic efficiency of the algal cells. The light that would normally reach the photosynthetic centres might be blocked by the adsorbed particles leached from mask. This would affect the rate of electron transport as well as the photosynthetic activity. One of the earlier study performed a CO_2_ depletion experiment and revealed that adsorbed plastic beads inhibited photosynthesis in *Scenedesmus obliquus* and *Chlorella sp.* ^49^. Our study also showed similar results. When the cell membrane becomes more permeable, both nTiO_2_ and FML can be internalized easily. This would exacerbate the damage to the cell’s structure and organelles like chloroplasts and mitochondria. Chlorophyll production and PSII may be inhibited if the chloroplast is damaged by nTiO_2_ and FML ^50^. On the other hand, HML caused a very minor decrease in PSII and Φm. This was because the concentration of the heavy metals was less, and so they were unable to permeate the algal cell membrane and could not damage the photosynthetic centres ^51^.

### 3.7. Surface chemical changes

The FTIR spectra of *Scenedesmus obliquus* treated with nTiO_2_, FML, and their binary combination are shown in Figure S6. The spectrum alterations that were seen on the cells of algae as a result of their interaction with the pollutants are summarized in Table 1. In the cells that were treated with individual nTiO_2_ and binary mixtures (nTiO_2_ + FML), the existence of Ti-O-Ti was verified by the appearance of a distinctive peak at 850 cm^-1^. Amide groups may be identified by their characteristic C=O and C-O stretching at 572 and 1020 cm^-1^ respectively. There are also two different peaks located at 1156 and 1246 cm^-1^, which represent C-O-C stretching and protein amides, respectively. Both 1450 and 1642 cm^-1^ represent CH_3_, CH_2_ deformations, and C-O stretching respectively. All of the samples that had been treated with FML revealed a peak at 2916 cm^-1^ that was connected with the asymmetric vibrations of nucleic acid. The peaks that correspond to –NH and –OH stretching vibrations may be seen at 3280 cm^-1^ in all of the treated samples.

**Table 1:** Spectral changes upon treatment with nTiO_2_, FML and nTiO_2_ + FML. ↑: Increased change in % Transmittance; ↑↑: Strongly increased change in % Transmittance; ↓: Decreased change in % Transmittance; ↓↓: Strongly change in % Transmittance

| Wavenumber<br>(cm<sup>-1</sup>) | Pristine<br>nTiO<sub>2</sub> treated | Pristine<br>FML treated | nTiO<sub>2</sub> +<br>FML treated | Peak assignment |
| --- | --- | --- | --- | --- |
| 572 | ↓ | ↑↑ | ↑ | C=O; out of the plane<br>in protein amides |
| 850 | ↑↑ | ↓ | ↑ | Attributed to the<br>presence of Ti-O-Ti |
| 1020 | ↑ | ↑↑ | ↑ | C-O stretching |
| 1156 | ↓ | ↑↑ | ↑ | C-O-C stretching |
| 1246 | ↓ | ↑ | ↑ | Protein amide III, α-<br>helix structure |
| 1450 | ↓↓ | ↑ | ↑ | CH<sub>3</sub> and CH<sub>2</sub><br>deformations |
| 1642 | ↑ | ↑ | ↑ | C-O stretching |
| 2916 | ↓↓ | ↑ | ↑ | Asymmetric vibrations<br>of nucleic acid |
| 3280 | ↑ | ↑ | ↑ | -NH and -OH bond<br>stretching of lipids |

According to the FTIR findings, one of the key factors that may have contributed to the toxicity of the binary combination is the interfacial interactions that occurred between the components of the algal cell wall and nTiO_2_ / FML. Microplastics in FML might form hetero-aggregates with nTiO_2_ and be adsorbed onto the algal cells. This causes damage to the algal cell membrane and can promote the internalization of nTiO_2_ ^18^. With low mixture concentrations, toxicity was also observed which might be due to the outcome of competition for binding sites to gain cellular entry ^52^.

## 3 Conclusions

The findings of this study demonstrated that disposable surgical face masks can be a potential source of microplastic as well as heavy metals to the environment. The freshwater ecosystem contains a variety of toxic nanoparticles and other natural substances. These microplastic leachates may combine with other particles and increase their toxicity towards algae. In this work, we have replicated this phenomenon by combining the whole face mask leachate with nTiO_2_ and observed its toxicity. When the mixture was exposed to *Scenedesmus obliquus*, it exhibited reduced cell growth and photosynthetic activity. Additionally, both oxidative stress and antioxidant activity were shown to be elevated. More research is needed to determine the impact of mask leachates along with other nanoparticles on the health of algae-eating freshwater species and whether or not it would travel up to higher trophic levels.

## Author Contributions

**Soupam Das:** Investigation, Methodology, Visualization, Formal analysis, Writing-original draft.

**Amitava Mukherjee:** Conceptualization, Methodology, Supervision, Project administration, Writing-Review and editing.

## Acknowledgment

The authors are thankful to Vellore Institute of Technology, Vellore, India for Field Emission Scanning electron microscope (FE-SEM) facilities used in this study. The authors would also like to acknowledge Indian Institute of Technology, Madras, India for inductively coupled plasma - optical emission spectrometry (ICP-OES) facility used in this study.

